# Insula dynorphin and kappa opioid receptor systems regulate alcohol drinking in a sex-specific manner in mice

**DOI:** 10.1101/2021.11.09.467893

**Authors:** Melanie M. Pina, Dipanwita Pati, Sofia Neira, Lisa R. Taxier, Christina M. Stanhope, Alexandra A. Mahoney, Shannon D’Ambrosio, Thomas L. Kash, Montserrat Navarro

## Abstract

Alcohol use disorder is complex and multi-faceted, involving the coordination of multiple signaling systems across numerous brain regions. Previous work has indicated that both the insular cortex and dynorphin (DYN)/Kappa opioid receptor (KOR) systems contribute to excessive alcohol use. More recently, we identified a microcircuit in the medial aspect of the insular cortex that signals through DYN/KOR. Here, we explored the role of insula DYN/KOR circuit components on alcohol intake in a long-term intermittent access (IA) procedure. Using a combination of conditional knockout strategies and site-directed pharmacology, we discovered distinct and sex-specific roles for insula DYN and KOR in alcohol drinking and related behavior. Our findings show that insula DYN deletion blocked escalated consumption and decreased overall intake of and preference for alcohol in male and female mice. This effect was specific to alcohol in male mice, as DYN deletion did not impact sucrose intake. Further, insula KOR antagonism reduced alcohol intake and preference during the early phase of IA in male mice only. Alcohol consumption was not affected by insula KOR knockout in either sex. In addition, we found that long-term IA decreased the intrinsic excitability of DYN and deep layer pyramidal neurons (DLPN) in the insula of male mice. Excitatory synaptic transmission was also impacted by IA, as it drove an increase in excitatory synaptic drive in both DYN neurons and DLPN. Combined, our findings suggest there is a dynamic interplay between excessive alcohol consumption and insula DYN/KOR microcircuitry.

**Significance Statement:** The insular cortex is a complex region that serves as an integratory hub for sensory inputs. In our previous work, we identified a microcircuit in the insula that signals through the kappa opioid receptor (KOR) and its endogenous ligand dynorphin (DYN). Both the insula and DYN/KOR systems have been implicated in excessive alcohol use and alcohol use disorder (AUD). Here, we utilize converging approaches to determine how insula DYN/KOR microcircuit components contribute to escalated alcohol consumption. Our findings show that insula DYN/KOR systems regulate distinct phases of alcohol consumption in a sex-specific manner, which may contribute to the progression to AUD.

## Introduction

Excessive alcohol consumption has a broad impact on human health, generating high personal and economic costs (Sacks et al., 2015). As a leading preventable cause of death in the United States, a large body of work has been devoted to identifying the brain structures and signaling mechanisms that drive excessive drinking and contribute to the development of alcohol use disorder (AUD). Preclinical research has shown the kappa opioid receptor (KOR) and its endogenous ligand dynorphin (DYN) are one of several key neurochemical systems involved in AUD (Crowley and Kash, 2015). For example, repeat variants in the genes that encode for KOR and DYN have been associated with an increased risk of AUD (Xuei et al., 2006; Edenberg et al., 2008; Karpyak et al., 2013). The involvement of DYN/KOR signaling in alcohol consumption and dependence has been further indicated by studies in rats showing that KOR antagonism increases alcohol self-administration (Mitchell et al., 2005) while agonism decreases volitional intake (Lindholm et al., 2001). Interestingly, the effects of KOR appear to reverse following alcohol dependence, as KOR antagonism can reduce excessive alcohol self-administration in post-dependent rats only (Walker and Koob, 2008; Walker et al., 2011). This finding suggests that chronic alcohol exposure shifts KOR signaling dynamics in a manner that drastically alters its function in alcohol intake.

As a multisensory hub, the insula plays a critical role in integrating internal and external stimuli to generate complex internal states and subsequent actions (Gehrlach et al., 2020). Connectivity patterns of the insula are extensive, with inputs and outputs to this region spanning across cortical and subcortical domains. Among the most predominate inputs to the insula, specifically to the medial and posterior portions, are the amygdala, thalamus, and sensory cortex, (Gehrlach et al., 2020). Notably, the reciprocal connectivity between the amygdala and insula has been well documented (e.g., (Allen et al., 1991; Augustine, 1996; McDonald et al., 1999; Santiago and Shammah-Lagnado, 2005) and this connection is believed to underlie the insula’s role in tastant reinforcement and gustatory valence encoding (Lavi et al., 2018; Schiff et al., 2018; Wang et al., 2018).

Human imaging studies have also implicated the insula as a region altered in AUD (Claus et al., 2011; Ihssen et al., 2011; Grodin et al., 2017), with findings supported by multiple rodent studies (Seif et al., 2013; Jaramillo et al., 2018a; Centanni et al., 2019; Chen and Lasek, 2020; Marino et al., 2021). In humans, PET imaging has shown there is reduced KOR availability in the insula of patients with AUD compared to controls (Vijay et al., 2018). Most notably, recent work has indicated that reductions in alcohol drinking and craving induced by the non-specific opioid antagonist naltrexone are, in part, associated with KOR availability in the insula (de Laat et al., 2019). Combined, these findings strongly suggest that the insula may serve as a critical locus for DYN/KOR action in AUD.

Beyond this work, there have been no comprehensive explorations of how DYN/KOR signaling in the insula can directly impact alcohol consumption. Here, we explored the role of insula DYN/KOR systems in both male and female mice using converging genetic and pharmacological approaches. In addition, we assessed how long-term alcohol consumption can impact insula neuronal function using slice physiology.

## Methods

### Mice

Male and female mice, aged 9-16 weeks at the start of the procedures, were singly housed and maintained on a reverse light cycle (12:12 hr light-dark) in a temperature-controlled colony room. Throughout the experimental procedures, mice were provided continuous access to food and water. In addition to Pdyn^IRES-Cre^ and Pdyn^IRES-Cre^ x Gt(ROSA26)Sor^loxSTOPlox-L10-GFP^(Pdyn^GFP^) (Krashes et al., 2014) mice, Pdyn^lox/lox^ (Bloodgood et al., 2021) and Oprk1^lox/lox^ (Crowley et al., 2016) conditional-knockout mice were generated as previously described and bred in-house. Wild-type C57BL/6J mice were purchased from The Jackson Laboratory. All experiments were performed in accordance with the NIH guidelines for animal research and with the approval of the Institutional Animal Care and Use Committee at the University of North Carolina at Chapel Hill.

### Alcohol and tastant drinking procedures

For drinking experiments, mice were singly housed and maintained on an Isopro RMH 3000 (LabDiet, St. Louis, MO, USA) diet, which has been shown to produce high levels of alcohol consumption (Marshall et al., 2015). To evaluate the effect of genetic knockout and KOR antagonism on alcohol consumption and preference, an intermittent access to alcohol (IA) procedure was run as previously described (Bloodgood et al., 2021). Briefly, mice were given 24 h home-cage access to a bottle containing a 20% w/v alcohol solution alongside a water bottle. Alcohol bottles were introduced 3 h into the dark cycle and bottles were weighed at the start and end of each 24 h session. Drinking sessions occurred three times per week with no less than 24 h and no more than 48 h between sessions (e.g., Mon, Wed, Fri). The amount of fluid lost due to passive drip was determined by placing alcohol and water bottles in an empty cage. Drip values were then calculated for each solution and total consumption was normalized to these values. To prevent the development of a side bias, the placement of alcohol bottles (left or right) was counterbalanced across sessions. To determine the selectivity of treatment effects on alcohol consumption, mice were given a sucrose challenge where they received 2 h access to a bottle of 3% sucrose in addition to water. Sucrose drinking experiments occurred 48 h after the final IA session.

### Elevated plus maze

To assay anxiety-related behavior, an elevated plus maze (EPM) task was used. The EPM apparatus consisted of two open (75 × 7 cm) and two closed (75 × 7 × 25 cm) arms arranged in a plus configuration wherein the arms of each type (open or closed) are positioned opposite of one another and connected via a central open zone (7 × 7 × 25 cm). During EPM testing, ambient illumination was maintained at around 15 lux. Mice were placed in the center zone of the apparatus at task onset, then allowed to explore freely for 5 minutes. Video recordings were obtained and analyzed by a blinded observer and with Ethovision 9.0 (Noldus Information Technologies) to determine the time spent in the open/closed arms and the total number of open arm entries.

### Surgical procedure

Mice were anesthetized by injection (i.p., 1.5 mL/kg) of ketamine/xylazine (Ketaset, code EA2489-564; AnaSed, NDC code 59399-111-50), then secured in a stereotaxic frame (Kopf Instruments) for intracranial viral and drug infusions. The insula (from bregma in mm: AP +0.86, ML ±3.59, DV −3.9) was targeted using standard mouse brain atlas coordinates (Franklin and Paxinos, 2008) and substances were microinjected using a 1 µl Neuros Syringe (33-gauge needle, Hamilton) controlled by an infusion pump. For conditional deletion, 300 nL/side of AAV5-CAMKIIα-Cre-eGFP (UNC vector core; 2.3 x 10^13^ vg/mL) or 200 nL/side AAV5-CMV-HI-eGFP-Cre-WPRE-SV4 (Addgene; ≥ 1 x 10^13^ vg/mL) was injected at a rate of 100 nL/min into the insula of Pdyn^lox/lox^ and Oprk1^lox/lox^ mice, respectively. After infusion, injectors were left in place for 5 min to allow for viral diffusion. To pharmacologically block KOR, nor-BNI (5 µg/µL, .5 µL/side in PBS) was infused into the insula of C57BL/6J mice over a period of 5 min. In a subset of mice, a GFP-tagged virus (AAV5-CMV-eGFP; 50 nL/side) was added to the nor-BNI mix to assess infusion spread. To minimize postoperative discomfort, meloxicam (5 mg/kg, i.p.; Metacam, NDC code 0010-6013) was administered at the time of surgery.

### Histology

Mice were deeply anesthetized with Avertin (250 mg/kg, IP) and transcardially perfused with chilled 0.01 M phosphate buffered saline (PBS, pH 7.4) followed by 4% paraformaldehyde (PFA). Brains were removed and immersed in 4% PFA overnight, then stored in 30% sucrose/PBS before 45 µm coronal sections were taken on a vibratome (Leica VT1000 S). Free-floating sections were processed for immunofluorescence to amplify GFP signal. Briefly, IC-containing sections (from bregma in mm, AP+1.54 to +0.38) were washed in PBS, then blocked and permeabilized in 5% normal donkey serum/0.3% Triton X-100/PBS for 45 min. Tissue was incubated overnight with gentle agitation at 4°C with a chicken polyclonal anti-GFP antibody (1:2000, Aves Labs) in blocking solution. Sections were rinsed, then blocked for 45 min before 2 h incubation at room temperature in Alexa Fluor 488-conjugated donkey anti-chicken IgG (1:400 in blocking solution, Jackson ImmunoResearch). Sections were rinsed in PBS after the final incubation and mounted with Vectashield Hardset Mounting Medium with DAPI (Vector Labs). Slides were imaged on an Olympus BX43 with attached optiMOS sCMOS camera (QImaging) or Keyence BZ-X800.

### Fluorescent in situ hybridization (FISH)

To validate Pdyn and Oprk1 deletion and compare basal levels of these transcripts between sexes, mice were anesthetized with isoflurane, rapidly decapitated, and brains rapidly extracted and snap frozen. Brains were then stored in a −80°C freezer prior to slicing. Sections (12 µm) were obtained using a Leica CM3050 S cryostat (Leica Microsystems, Wetzlar, Germany), mounted onto Superfrost Plus slides (Fisher Scientific, Hampton, NH), and stored at −80°C prior to in situ hybridization. RNAscope was performed according to the manufacturer’s instructions (Advanced Cell Diagnostics, Newark, CA). Briefly, sections were fixed using 4% PFA, dehydrated, and washed with protease IV solution (Advanced Cell Diagnostics, Newark, CA) prior to incubation with probes for Pdyn and Oprk1 (Advanced Cell Diagnostics, Newark, CA). Slides were then coverslipped using ProLong Gold Antifade Mountant with DAPI, and stored at −80°C. Images were captured using a Zeiss 800 upright confocal microscope (Hooker Imaging Core, UNC-Chapel Hill). For examination of basal sex differences in levels of Pdyn, Oprk1, and Vgat mRNA, tiled z-stacks of bilateral insula were captured at 20x magnification (3 slices/animal), and maximum projection intensity images were generated using Zen blue software (Zeiss). Analysis was performed using subcellular particle detection in QuPath (Bankhead et al, 2017). Cells were considered Pdyn, Oprk1, and/or Vgat-positive if they contained ≥ 5 puncta in each probe’s respective channel. For examination of Pdyn and Oprk1 knockdown, validation of AAV-Cre injection into the insula was performed by confirming the presence of GFP at the target site, and only sections with GFP were used for imaging and subsequent analysis. Tiled z-stacks of bilateral insula (2 slices/mouse) were captured and converted to maximum projection intensity images as described above. Mean fluorescence intensity/mm^2^ was calculated using QuPath (Bankhead et al., 2017).

### Electrophysiology Recordings

Whole-cell patch-clamp recordings were obtained from the insula of Pdyn^GFP^ mice. After rapid decapitation under isoflurane anesthesia, brains were quickly extracted and immersed in a chilled and carbogen (95% O_2_/5% CO_2_)-saturated sucrose artificial cerebrospinal fluid (aCSF) cutting solution (in mM): 194 sucrose, 20 NaCl, 4.4 KCl, 2 CaCl_2_, 1 MgCl_2_, 1.2 NaH_2_PO_4_, 10 D-glucose and 26 NaHCO_3_. Coronal slices (300 µM) containing the insula were prepared on a vibratome, then transferred to a holding chamber containing heated oxygenated aCSF (in mM: 124 NaCl, 4.4 KCl, 1 NaH_2_PO_4_, 1.2 MgSO_4_, 10 D-glucose, 2 CaCl_2_, and 26 NaHCO_3_). After equilibration (≥ 30 min), slices were placed in a submerged recording chamber superfused (2 mL/min) with oxygenated aCSF warmed to approximately 30-35°C. Neurons were visualized under a 40x water immersion objective with video-enhanced differential interference contrast, and a mercury arc lamp-based system was used to visualize fluorescently labeled pDyn^GFP^ neurons. Recording pipettes (2–4 MΩ) were pulled from thin-walled borosilicate glass capillaries. Signals were acquired using an Axon Multiclamp 700B amplifier (Molecular Devices), digitized at 10 kHz, filtered at 3 kHz, and analyzed in pClamp 10.7 or Easy Electrophysiology. Series resistance (*R_a_*) was monitored without compensation and data were discarded from recordings where changes in *R_a_* exceeded 20%.

Intrinsic properties and action potentials were recorded using a potassium gluconate-based intracellular solution (in mM): 135 K-gluconate, 5 NaCl, 2 MgCl_2_, 10 HEPES, 0.6 EGTA, 4 Na_2_ATP, 0.4 Na_2_GTP, pH 7.3, 289–292mOsm. Input resistance and sag ratio were obtained in voltage clamp mode and sag ratio was obtained from a negative −100 mV current step and defined as the sag divided by the minimum voltage deflection from baseline. Current clamp recordings were obtained from insula Pdyn and deep layer pyramidal neurons at resting membrane potential. Current-injection evoked action potentials were evaluated by measuring the following: (1) action potential threshold (voltage at which cell first fired) and rheobase (minimum current required to evoke an action potential), both determined using a linearly increasing 120 pA/sec ramp protocol; and (2) the number of spikes fired at increasing current steps (50 pA increments, 0 to 650 pA). Action potential kinetics including (1) peak amplitude, (2) half-width, (3) rise time, and (4) decay time were obtained from the current step protocol and action potential thresholds were estimated using Method II (Sekerli et al., 2004) and analyzed using Easy Electrophysiology.

Spontaneous synaptic events were recorded in voltage-clamp mode with a cesium methanesulfonate-based intracellular solution (in mM): 135 cesium methanesulfonate, 10 KCl, 1 MgCl_2_, 10 HEPES, 0.2 EGTA, 4 MgATP, 0.3 Na_2_GTP, 20 phosphocreatine, pH 7.3, 285-290 mOsm with 1mg/mL QX-314. The ratio of excitatory to inhibitory (E:I ratio) transmission was measured by isolating glutamate (*V_hold_* = −55 mV) and GABA currents (*V_hold_* = +10 mV) within individual neurons.

### Experimental Design and Statistical Analysis

Data were analyzed by mixed model three-way (sex x group x session/time; sex x treatment x time/current) or two-way (sex x group/treatment; group/treatment x session/time/current) ANOVA, where group = GFP/Cre or Vehicle/nor-BNI and treatment = H2O/IA. Additional analyses were performed using one-way ANOVA or t-tests to examine the effects of group and sex on Pdyn and Oprk1 transcript levels, alcohol intake/preference, total fluid intake, sucrose intake, time spent in open/closed arms of EPM or the effects of treatment on RMP, rheobase, sE/IPSC frequency/amplitude, and E/I ratio. Where data violated assumptions of normality or homoscedasticity, a Mann-Whitney U test or Welch’s correction was used, respectively, for analysis. Data are presented as mean ± SEM and were analyzed using JASP 0.14 (JASP Team) or in Prism 9 (GraphPad Software, LLC). Significance threshold was set at α = 0.05, and post-hoc pairwise comparisons were multiplicity adjusted.

## Results

### Knockout of Pdyn in the insula decreases alcohol drinking in male and female mice

To define the role of insula Dyn neurons in alcohol intake, we conditionally deleted Pdyn in male and female mice prior to initiation of an 8-week IA drinking procedure (Fig. 1A). Anxiety-like behavior was also assessed in the EPM test 24h following the final IA session. To assess the specificity of knockout on alcohol intake, mice were given a 2h sucrose drinking challenge 24h after EPM testing. Site-specific knockout of Pdyn was achieved by infusing AAV5 with CaMKIIa-promoter driven expression of a GFP-fused Cre recombinase (n = 8 male / 9 female) or a GFP only control (n = 6 male / 7 female) into the insula of Pdyn^lox/lox^ mice. Figure 1B shows overlay of viral expression mapped at + 0.98 mm (from bregma) for male and female GFP (blue) and Cre (magenta) group mice. Representative photomicrographs of GFP expression and Pdyn mRNA expression in the insula of GFP (top) and Cre (bottom) mice are shown in Figs. 1C and 1D, respectively. In situ hybridization showed that compared to GFP controls, injection of a Cre encoding vector led to a significant reduction in insula Pdyn mRNA expression as indexed by mean fluorescent intensity [n = 2 mice/sex, *t_6_* = 3.33, *p* = 0.016] (Fig. 1E). Additionally, we found no difference in basal Pdyn mRNA expression in naïve male or female mice [n = 3/sex, *t_4_* = 1.60, *p* = 0.186] (Fig. 1F). Knockout of Pdyn decreased alcohol intake and preference in males (Fig. 1G-H) [group x session interaction: *F*_(23,276)_ = 1.97, p = 0.006, intake / *F*_(23,276)_ = 2.62, *p* < 0.001, preference; group main effect: *F*_(1,12)_ = 4.62, *p* = 0.053, intake / *F*_(1,12)_ = 6.59, *p* = 0.025, preference; session main effect: *F*_(23,276)_ = 3.89 *p* < 0.001, intake / *F*_(23,276)_ = 3.96, *p* < 0.001, preference] and females (Fig. 1I-J) [group x session interaction: *F*_(23,322)_ = 1.61, *p* = 0.039, intake / *F*_(23,322)_ = 1.33, *p* = 0.143, preference; group main effect: *F*_(1,14)_ = 11.5, *p* = 0.004, intake / *F*_(1,14)_ = 9.39, *p* = 0.008, preference; session main effect: *F*_(23,322)_ = 9.33, *p* < 0.001, intake / *F*_(23,322)_ = 7.92, *p* < 0.001, preference]. Knockout of Pdyn in the insula also blocked escalation of alcohol intake (Fig. 1K, N), as average intake levels significantly increased from weeks 1-4 to weeks 5-8 in controls (GFP) [Males: *t*_5_ = 4.42, *p* = 0.007; Females: *t*_6_ = 3.56, *p* = 0.012] but not in Pdyn knockout (Cre) mice [Males: *p* = 0.378; Females: *p* = 0.083].

**Figure 1.**
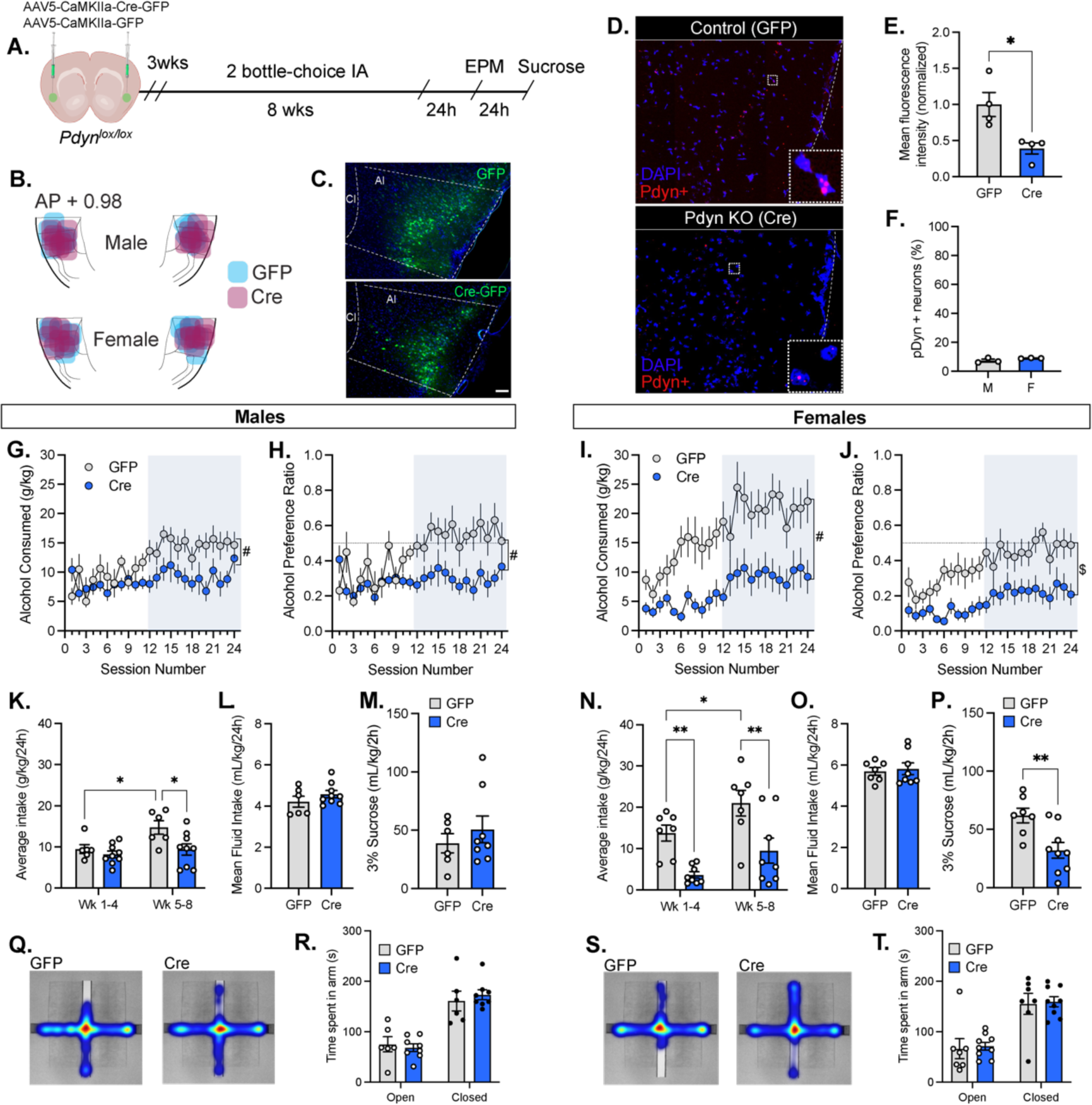
Genetic deletion of insula Pdyn blocks intermittency-induced escalation of alcohol intake in male and female mice. (A) Experimental timeline: Following AAV injections and a 3 wk delay for expression, male and female Pdyn floxed mice were run through an 8-wk 2-bottle choice intermittent access to alcohol (IA) procedure. 24h after the final alcohol exposure, mice were tested for anxiety-like behavior in the elevated plus maze (EPM), then given a 2-h sucrose challenge 24h thereafter. (B) Map of viral spread at +0.98 mm (from bregma) in male and female mice in GFP (blue) and Cre (magenta) groups. (C) Representative images showing expression following injection of AAV-CAMKIIa-GFP control vector (top) and AAV-CAMKIIa-Cre-GFP (bottom) in the mouse insula. Cl, claustrum; AI, agranular insula; scale bars = 100 uM. (D-F) In situ hybridization was used for validation of Pdyn knockout and comparisons of basal Pdyn mRNA expression in male and female mice. (D) Representative images of Pdyn mRNA expression in a GFP (control) mouse and a Cre (knockout) mouse. (E) Compared to GFP controls, mice injected with an an AAV encoding for Cre recombinase showed a significant reduction in Pdyn mRNA expression in the insula, as indexed by mean fluorescence intensity. (F) In naïve male and female mice, there was no difference in basal Pdyn mRNA expression. (G-J) Insula Pdyn knockout significantly reduced alcohol consumption and preference in (G-H) male and (I-J) female mice. This effect was most prominent during the final 4 wks of drinking (highlighted blue), during which (K) male and (F) female GFP control mice consumed significantly more alcohol as compared to the first 4 wks of IA. Cre mice did not exhibit this escalated pattern of intake and showed lower average intake levels compared to GFP controls, with (K) males consuming significantly less alcohol at weeks 5-8 and (N) females at wks 1-4 and 5-8. (L-M) In male mice, the effect of Pdyn deletion was specific to alcohol, as it did not affect mean fluid intake (L) or intake of a 3% sucrose solution (M). (O-P) In female mice, Pdyn deletion did not impact (O) mean fluid intake while it (P) decreased sucrose consumption. (Q-T) There was no effect of insula Pdyn knockout on anxiety-like behavior as measured in the EPM test. (Q, S) Representative heat maps showing time spent in EPM arms in (Q) male and (S) female mice in GFP (left panel) and Cre (right panel) groups. In both (R) male and (S) female mice, Cre and GFP groups spent a similar amount of time in the open and closed arms of the EPM. # group x sex interaction, p < 0.05; $ group main effect, p < 0.01; * p < 0.05; ** p < 0.01; *** p < 0.001

Total fluid intake was not affected by Pdyn knockout in male [*p* = 0.249] or female mice [*p* = 0.737] (Fig. 1L, O). Interestingly, Pdyn deletion decreased sucrose consumption in females [*t*_14_ = 3.17, *p* = 0.007] (Fig. 1P), but this effect was absent in male mice [*p* = 0.737] (Fig. 1M). Combined, these findings show that Pdyn deletion in the insula selectively reduces alcohol intake in male mice and suggest that insula Pdyn may play a key role in driving escalated alcohol consumption. We next tested the impact of Pdyn deletion on anxiety-like behavior by exposing mice to an EPM test 24h following their final drinking session (Fig. 1Q-T). We found no effect of Pdyn knockout in either sex on time spent in the open [Males: *p* = 0.671; Females: *p* = 0.252] or closed [Males: *p* = 0.188; Females: *p* = 0.843] arms of the EPM 24h following their final drinking session (Fig. 1R, T).

### Local infusion of nor-BNI decreases early phase alcohol drinking in male, but not female mice

Previous studies have found that the KOR antagonist nor-BNI can reduce alcohol consumption when given systemically (Walker et al., 2011; Anderson et al., 2016). To assess insula KOR involvement in alcohol drinking, we locally infused the long-acting antagonist nor-BNI (Munro et al., 2012) or PBS vehicle into male (n = 12 nor-BNI / 10 PBS) and female (n = 10 nor-BNI / 10 PBS) C57BL/6J mice prior to the start of an 8-week IA procedure (Fig. 2A-B). To determine the spread of nor-BNI infusion within the insula, a subset of female mice were co-injected with AAV5-CMV-eGFP (50 nL/side). Representative images showing GFP expression and cannula tracts along with reconstructed maps of injector tip placements are shown in Fig. S2. Previous findings show that a single injection of nor-BNI blocks KOR for up to 3 weeks (Horan et al., 1992; Bruchas et al., 2007). Thus, the prolonged action of nor-BNI allowed us to pharmacologically assess the role of insula KOR in alcohol consumption over multiple weeks. As above, mice were also assessed for anxiety-like behavior (EPM) and sucrose intake at 24h and 48h, respectively, following the final drinking session. As shown in Fig. 2B, female mice were given one additional assay, a saccharine challenge, 24h after the sucrose challenge.

**Figure 2.**
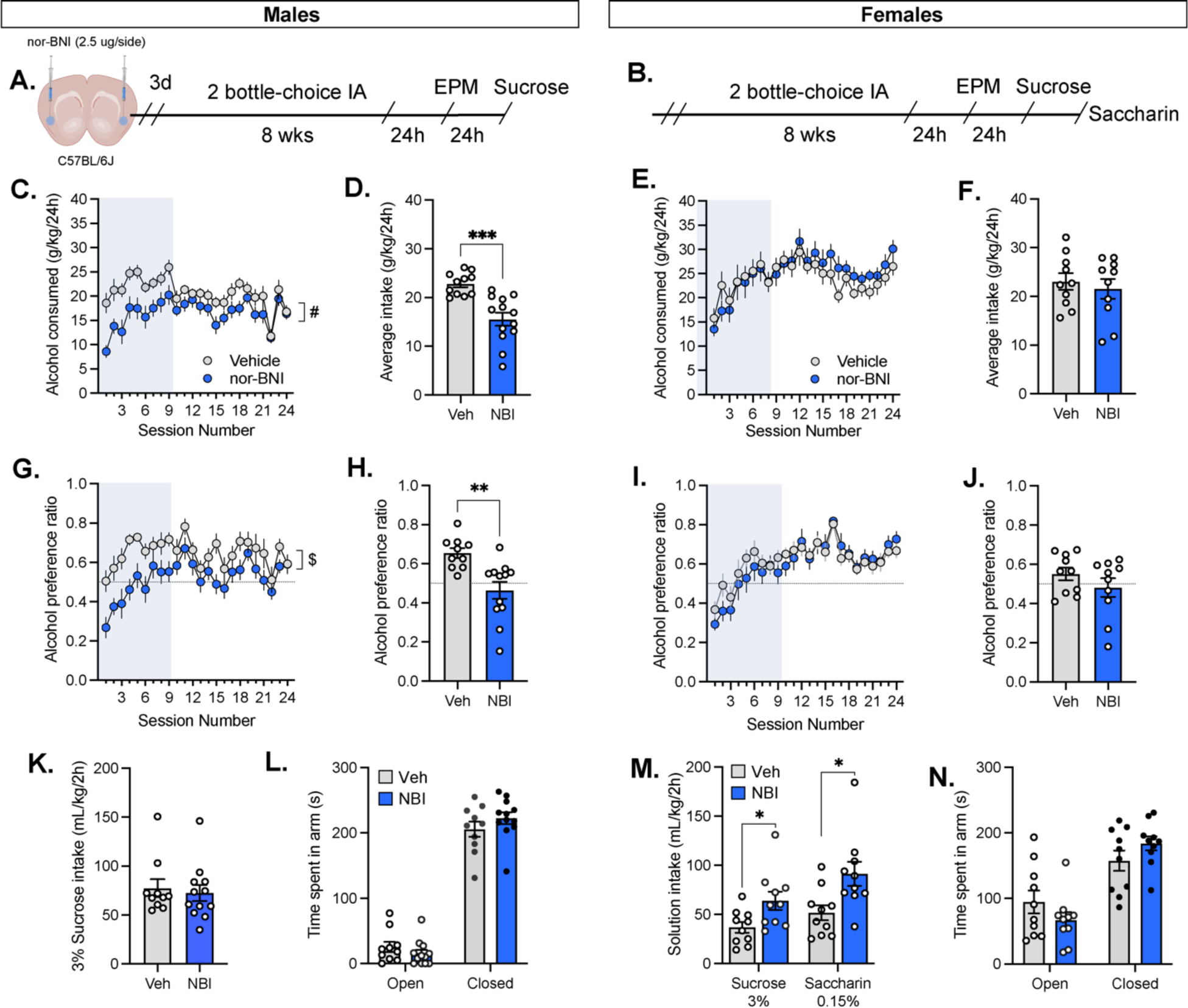
Pharmacological blockade of insula KOR by nor-BNI decreases alcohol intake in early phases of drinking in male mice and increases intake of sweet solutions in female mice. (A-B) Experimental timeline: The KOR antagonist nor-BNI (2.5 ug/side) or PBS (vehicle) was microinjected into the insula of C57BL/6J (A) male and (B) female mice about 3 days before exposure to an 8-wk 2-bottle choice intermittent access to alcohol (IA) procedure. 24h after their final alcohol exposure, mice were tested for anxiety-like behavior in the elevated plus maze (EPM), then given a 2h sucrose challenge 24h later, and female mice received a saccharine challenge 24h thereafter. In (C, G) male but not (E, I) female mice, nor-BNI (NBI) produced a transient decrease in alcohol consumption (C, E) and preference (G, I) during the first 3 weeks or 9 drinking sessions (highlighted blue). (D, H) During the first 9 sessions of IA in male mice, average alcohol intake (D) and preference (H) levels were significantly lower in NBI mice compared to vehicle (Veh) controls. (F, J) In female mice, there was no impact of NBI on alcohol (F) consumption and (J) preference during the first 9 wks of IA. (K) There was no effect of NBI on sucrose intake in males, suggesting KOR antagonism selectively reduced the consumption of alcohol. (M) In female mice, NBI increase both sucrose and saccharine intake compared to Veh. (L, N) NBI did not impact EPM performance in either sex. # group x sex interaction, p < 0.05; $ group main effect, p < 0.01; * p < 0.05; ** p < 0.01; *** p < 0.001.

Our findings show that in male mice, insula KOR antagonism by nor-BNI reduced alcohol consumption (Fig. 2C) [group x session: *F*_(23,460)_ = 2.08, *p* = 0.003; main effects of group: *F*_(1,20)_ = 8.27, *p* = 0.009; and session: *F*_(23,460)_ = 8.07, *p* < 0.001] and alcohol preference (Fig. 2G) [group x session: *F*_(23,460)_ = 1.45, *p* = 0.082; main effects of group: *F*_(1,20)_ = 6.98, *p* = 0.016; and session: *F*_(23,460)_ = 6.93, *p* < 0.001]. Conversely, in female mice, we found no effect of KOR antagonism on intake (Fig. 2E) [group x session: *p* = 0.281; group main effect: *p* = 0.647; session main effect *F*_(23,414)_ = 8.78, *p* < 0.001] or preference (Fig. 2I) [group x session: *p* = 0.378; group main effect: *p* = 0.692; session main effect [*F*_(23,414)_ = 16.3, *p* < 0.001]. The reduction in intake by nor-BNI in males appeared to occur predominately during the first 9 sessions or 3 weeks of the IA procedure. Thus, we evaluated this by collapsing alcohol intake over the first 9 sessions of IA and found that nor-BNI decreased average alcohol intake (Fig. 2D) [*t*_22_ = 4.56, *p* < 0.001] and alcohol preference (Fig. 2H) [*t*_20_=3.58, *p* = 0.002] in male mice but not female mice (Fig. 2F, J) [intake: *p* = 0.578; preference: *p* = 0.245]. Nor-BNI did not affect intake of a 3% sucrose solution in male mice (Fig. 2K) [*p* = 0.691], but, led to a significant increase in sucrose intake in female mice (Fig. 2M) [*t*_18_ = 2.52, *p* = 0.022]. As a follow-up in females, we tested whether nor-BNI would affect the consumption of saccharin, a non-caloric sweet solution. Female mice were given access to a 0.15% saccharin solution and intake was measured at the end of a 2-h session. As with sucrose, insula KOR antagonism increased saccharin intake, as female nor-BNI mice consumed significantly more of the sweetened solution than did vehicle controls and the non-caloric sweetener saccharine at a 0.15% concentration (Fig. 2M) [*t*_18_ = 2.73, *p* = 0.014]. Thus, while insula KOR antagonism can selectively decrease alcohol intake in males, it may drive increased consumption of palatable solutions in females. Finally, we tested anxiety-like behavior in the EPM and found that nor-BNI did not affect time spent in the open arms or closed arms of the apparatus in male mice (Fig. 2L) [open: *p* = 2.80; closed: *p* = 0.213] or in female mice (Fig. 2N) [open: *p* = 0.213; closed: *p* = 0.179].

In a follow-up experiment, we administered nor-BNI 7 weeks into drinking procedure to (1) test if the transient reduction in alcohol intake it produced in male mice was due to a waning effect of the drug and (2) whether nor-BNI would impact females at this later phase of intake. Here, male (n = 10 nor-BNI / 11 PBS) and female (n = 8 nor-BNI / 9 PBS) C57BL/6J mice underwent 7 weeks of IA to alcohol before nor-BNI was delivered locally into the insula (Fig. 3A). After recovery, mice were given 3 additional weeks (9 sessions) of IA followed by a 3% sucrose challenge. When the final 9 session of IA were assessed, we found there was no effect of the nor-BNI on alcohol intake or preference in males (Fig. 3B, D) [intake (Fig. 3B): group x session interaction: *p* = 0.594; group main effect: *p* = 0.803; session main effect: *F*_(8,152)_ = 4.86, *p* < 0.001; preference (Fig. 3D): group x session interaction: *p* = 0.356; group main effect: p = 0.564; session main effect: *F*_(8,152)_ = 2.41, *p* = 0.018] and females (Fig. 3F, H) [intake (Fig. 3F): group x session interaction: *p* = 0.717; group main effect: *p* = 0.458; session main effect: *F*_(8,120)_ = 4.42, *p* < 0.001; preference (Fig. 3H): group x session interaction: *p* = 0.219; group main effect: p = 0.464; session main effect: *F*_(8,120)_ = 3.69, *p* < 0.001]. Similarly, when data were collapsed across the final 9 session of IA, we found no effect of insula nor-BNI infusion on average levels of alcohol intake and preference in males or females (*t* < 1, for all analyses).

**Figure 3.**
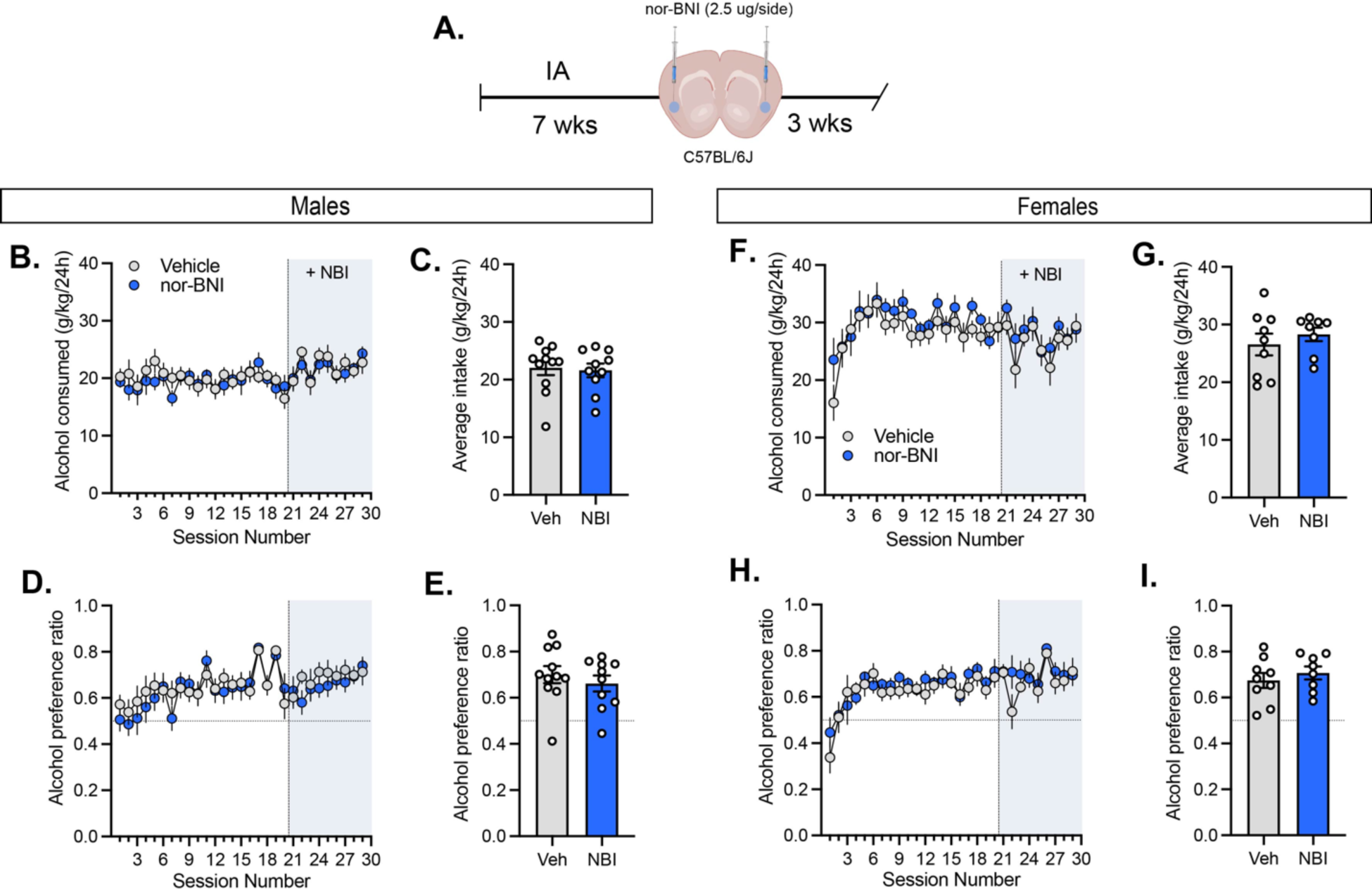
Pharmacological blockade of insula KOR by nor-BNI does not affect alcohol drinking in male or female mice during later stages of intake. (A) Experimental timeline: To test the effect of KOR antagonism on the later phase of alcohol drinking, mice were first exposed to 7 weeks of IA before receiving intra-insula injections nor-BNI (NBI, 2.5 ug/side) or PBS (vehicle). Following recovery, IA was resumed for an additional 3 wks. (B-E) In both male and (F-I) female mice, NBI did not affect (B, F) intake or (D, H) preference for alcohol when injected after 7 wks of IA. When the final 3 wks (8 sessions, post injection) were analyzed for (C, E) male and (G, I) female mice, there was no difference in average alcohol intake (C, G) or preference (E, I) between NBI and Veh groups.

### Knockout of KOR in the insula does not alter alcohol drinking in male or female mice

Our nor-BNI data suggest that in male mice, KOR localized within the insula can regulate aspects of alcohol consumption. However, KOR can be expressed at multiple sites within a given brain region, including on presynaptic nerve terminals. Given that nor-BNI can also act presynaptically at KOR-expressing afferent terminals, we wanted to more selectively target locally-expressed KOR in the insula. Using an *Oprk1^l^*^ox/lox^ mouse line, we selectively deleted KOR in the insula by injecting AAV5 encoding for a GFP-tagged Cre recombinase (n = 15 male / 16 female) or a GFP only control (n = 13 male / 15 female) (Fig. 4A). Following recovery, mice underwent 8 weeks of IA before being tested for anxiety-like behavior (EPM) and sucrose intake. Overlay of viral expression mapped at + 0.98 mm (from bregma) for male and female GFP (blue) and Cre (magenta) group mice is illustrated in Fig. 4B. Representative photomicrographs of viral expression and Oprk1 mRNA expression in the insula of GFP (top) and Cre (bottom) mice are shown in Figs. 4C and 4D, respectively. Using in situ hybridization, we found that viral-mediated delivery of Cre recombinase significantly reduced Oprk1 mRNA expression compared to a GFP control, as indexed by mean fluorescent intensity [n = 2-3 mice/sex, *t_9_* = 3.30, *p* = 0.009] (Fig. 4E). We found no difference in basal Oprk1 mRNA in naïve male or female mice [n = 3/sex, *t_4_* = 1.70, *p* = 0.164] (Fig. 4F), indicating that males and females express similar KOR transcript levels. In female mice, knockout of insula KOR had no effect on alcohol intake (Fig. 4I) [group x session interaction: *p* = 0.123; group main effect: *p* = 0.734; session main effect: *F*_(23,667)_ = 8.99, *p* < 0.001] or alcohol preference (Fig. 4J) [group x sex interaction: *p* = 0.331; group main effect, *p* = 0.519; session main effect: *F*_(23,667)_ = 9.27, *p* < 0.001] across the 8 week IA procedure. In male mice, analyses yielded a significant group x session interaction on intake (Fig. 4G) [*F*_(23,575)_ = 1.56, *p* = 0.047] in addition to a main effect of session [*F*_(23,575)_ = 2.79, *p* < 0.001] but not group [*p* = 0.097]. This finding may be due, in part, to the lack of escalation in GFP control mice and the increased intake in KOR knockout mice during the final week of IA (sessions 22-24). However, we found no effect of KOR deletion on alcohol preference in males (Fig. 4H) [group x session interaction: *p* = 0.484; group main effect, *p* = 0.162; session main effect: *F*_(23,575)_ = 2.33, *p* < 0.001]. To further assess intake and preference, we examined the first and latter halves (weeks 1-4 and 5-8) of the IA procedure. We found that from weeks 1-4 to weeks 5-8, female mice escalated their average alcohol intake (Fig. 4N) [paired t-tests; GFP: *t*_14_ = 3.98, *p* < 0.001; Cre: *t*_15_ = 4.42, *p* < 0.001] whereas male mice did not (Fig. 4K) [GFP: *p* = 0.298; Cre: *p* = 0.700]. In both males and females, KOR knockout did not affect total fluid intake [Males (Fig. 4L): *p* = 0.587; Females (Fig. 4O): *p* = 0.654] or 3% sucrose intake [Males (Fig. 4): *p* = 0.810; Females (Fig. 4P): *p* = 0.594]. Additionally, there was no difference in anxiety-like behavior between GFP and Cre groups, as indexed by time spent in the open and closed arms of the EPM apparatus in both males (Fig. 4R) [open: *p* = 0.318, closed: *p* = 0.117] and females (Fig. 4 T) [open: *p* = 0.711, closed: *p* = 0.654]. Representative heat map images illustrating time spent in the open and closed arms of the EPM for GFP (left) and Cre (right) male and female mice are shown in Figs. 4Q and 4S, respectively.

**Figure 4.**
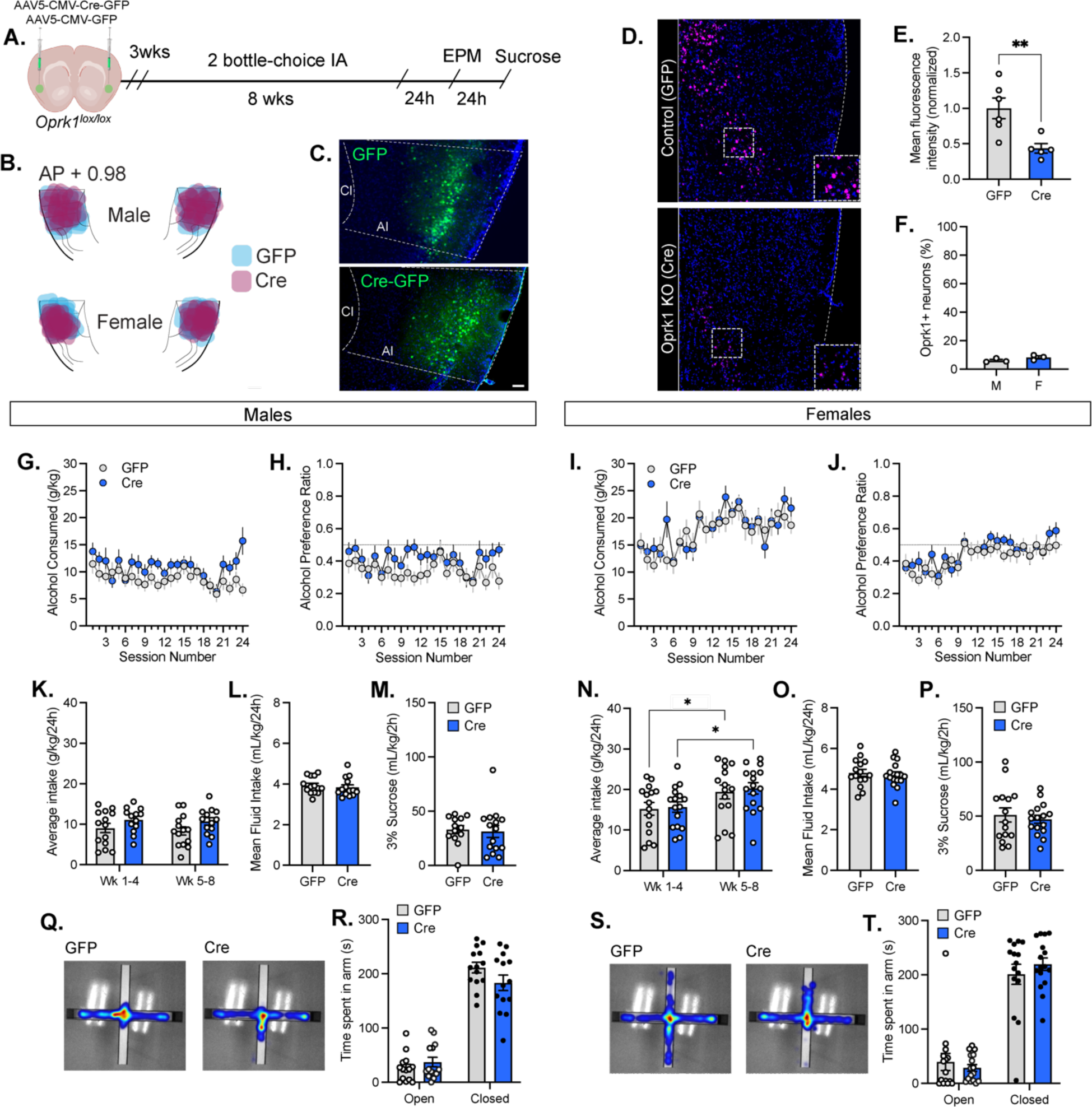
Genetic deletion of insula KOR does not impact alcohol intake in male or female mice. (A) Experimental timeline: 3 wks following AAV injections, male and female floxed Oprk1 mice were run through an 8-wk 2-bottle choice intermittent access to alcohol (IA) procedure. 24h after the final alcohol exposure, mice were tested for anxiety-like behavior in the elevated plus maze (EPM), then given a 2-h sucrose challenge 24h thereafter. (B) Map of viral spread at +0.98 mm (from bregma) in male and female mice in GFP (blue) and Cre (magenta) groups. (C) Representative images showing expression following injection of AAV-CMV-GFP control vector (top) and AAV-CMV-Cre-GFP (bottom) in the insula. Cl, claustrum; AI, agranular insula; scale bars = 100 uM. (D-F) In situ hybridization was used for validation of Oprk1 knockout and comparisons of basal Oprk1 mRNA expression in male and female mice. (D) Representative images of Oprk1 mRNA expression in a GFP (control; top) mouse and a Cre (knockout; bottom) mouse. (E) Compared to GFP controls, mice injected with an an AAV encoding for Cre recombinase showed a significant reduction in the number of Oprk1+ cells in the insula. (F) In naïve male and female mice, there was no difference in basal Oprk1 mRNA expression. (G-J) Compared to GFP mice, Cre-mediated deletion of insula KOR did not affect (G, I) alcohol consumption or (H, J) preference in (G-H) male or (I-J) female mice. (K-M) In males and (N-P) females, there was no difference in (K, N) alcohol intake, (L, O) mean fluid intake, or (M, P) sucrose intake (G) between GFP and Cre groups. (N) Female but not (K) male mice showed escalated alcohol intake between wks 1-4 and 5-8. (Q-T) There was no effect of insula KOR knockout on anxiety-like behavior as measured in the EPM test. (Q, S) Representative heat maps showing time spent in EPM arms in (Q) male and (S) female mice in GFP (left panel) and Cre (right panel) groups. In both (R) male and (S) female mice, Cre and GFP groups spent a similar amount of time in the open and closed arms of the EPM

### Long-term alcohol drinking exerts a sex-dependent effect on insula neuronal function

Previously, we identified a microcircuit in the medial agranular insular cortex that is modulated by KOR (Pina et al., 2020). Within this microcircuit, Pdyn neurons are densely clustered in layer 2/3 and KOR expression is localized to deep layer 5/6. Functionally, activation of insula KOR by Dyn dampens local inhibitory tone, which leads to a disinhibition of and an increase in excitatory synaptic drive in deep layer pyramidal neurons (DLPN). Thus, based on our above findings we wanted to determine the effect of long-term alcohol drinking on neuronal function in insula Pdyn and DLPN neurons. To assess the effect of long-term alcohol drinking on Pdyn neuronal function, adult male and female Pdyn^GFP^ mice were exposed to 8 weeks of IA to a 20% alcohol solution or a water control before tissue was collected for whole-cell patch clamp electrophysiology 24h after the final IA session (Fig. 5). We first assessed the intrinsic properties of Pdyn neurons in H2O controls and IA male and female mice, including resting membrane potential (RMP), input resistance (Rm), and sag ratio (Fig. 5A-C). We found a sex-dependent effect of IA on RMP (Fig. 5A), as indicated by a significant sex x treatment interaction [*F_(1,36)_* = 7.13, *p* = 0.011] but no effect of sex [p = 0.908] or treatment [*p* = 0.591]. Follow-up analyses revealed that IA significantly increased RMP in female mice [*U* = 23.5, *p* = 0.025] but not in male mice [*p* = 0.151]. In both male (n = 9 cells/3 mice EtOH, n = 10 cells/4 mice H2O) and female (n = 11 cells/5 mice EtOH, n = 10 cells/4 mice H2O) mice, IA did not alter input resistance (Fig. 5B) [treatment x sex interaction: *p* = 0.343; treatment main effect: *p* = 0.321] or sag ratio (Fig. 5C) [treatment x sex interaction: *p* = 0.147; treatment main effect: *p* = 0.599] of insula Pdyn neurons. There was no effect of sex on sag ratio (*p* = 0.192), however, in female mice Pdyn neurons exhibited a lower input resistance as indicated by a significant main effect of sex [F_(1, 36)_ = 6.72, *p* = 0.014]. We next assessed the impact of IA on Pdyn neuron excitability and found that 8 weeks of IA decreased the excitability of these neurons in male but not female mice (Fig. 5D-I). This was supported by a significant decrease in firing to increasing current steps (Fig. 5D) in males [treatment x current interaction: *F*_(10,170)_ = 7.75, *p* < 0.001; treatment main effect: *F*_(1,17)_ = 9.69, *p* = 0.006; current main effect: *F*_(10,170)_ = 40.90, *p* < 0.001] but not females [treatment x current interaction: *p* = 0.999; treatment main effect: *p* = 0.993; current main effect: *F*_(10,190)_ = 18.5, *p* < 0.001]. Additionally, the rheobase (minimum current to elicit firing) of Pdyn neurons was significantly higher in male IA mice compared to H2O controls (Fig. 5H) [*t*_15_ = 2.19, *p* = 0.042], and there was no effect of IA in female mice [*p* = 0.859]. There was no effect of IA on action potential (AP) threshold in either sex (Fig. 5G) [treatment x group interaction: *p* = 0.403; treatment main effect: *p* = 0.186, sex main effect: *p* = 0.549]. Representative traces show Pdyn neuron firing to a 600 pA current step (Fig. 5E) and a 120 pA/s ramp protocol (Fig. 5I) in male (left, blue) and female (right, purple) H2O and IA mice. Finally, we analyzed action potential kinetics and found no effects of IA on AP peak amplitude (Fig 5J) [Fs < 1 for treatment x sex interaction, treatment and sex main effects], half-width (Fig 5K) [Fs < 1 for treatment x sex interaction and treatment main effect, sex main effect, p = 0.058], rise (Fig 5L) [Fs < 1 for treatment x sex interaction and treatment main effect, sex main effect, p = 0.073], and decay (Fig 5M) [Fs < 1 for treatment x sex interaction and treatment main effect, sex main effect, p = 0.037]. Thus, the only significant effect found on Pdyn neuron AP kinetics was a main effect of sex on AP decay time, as female mice exhibited faster decay times as compared to males.

**Figure 5.**
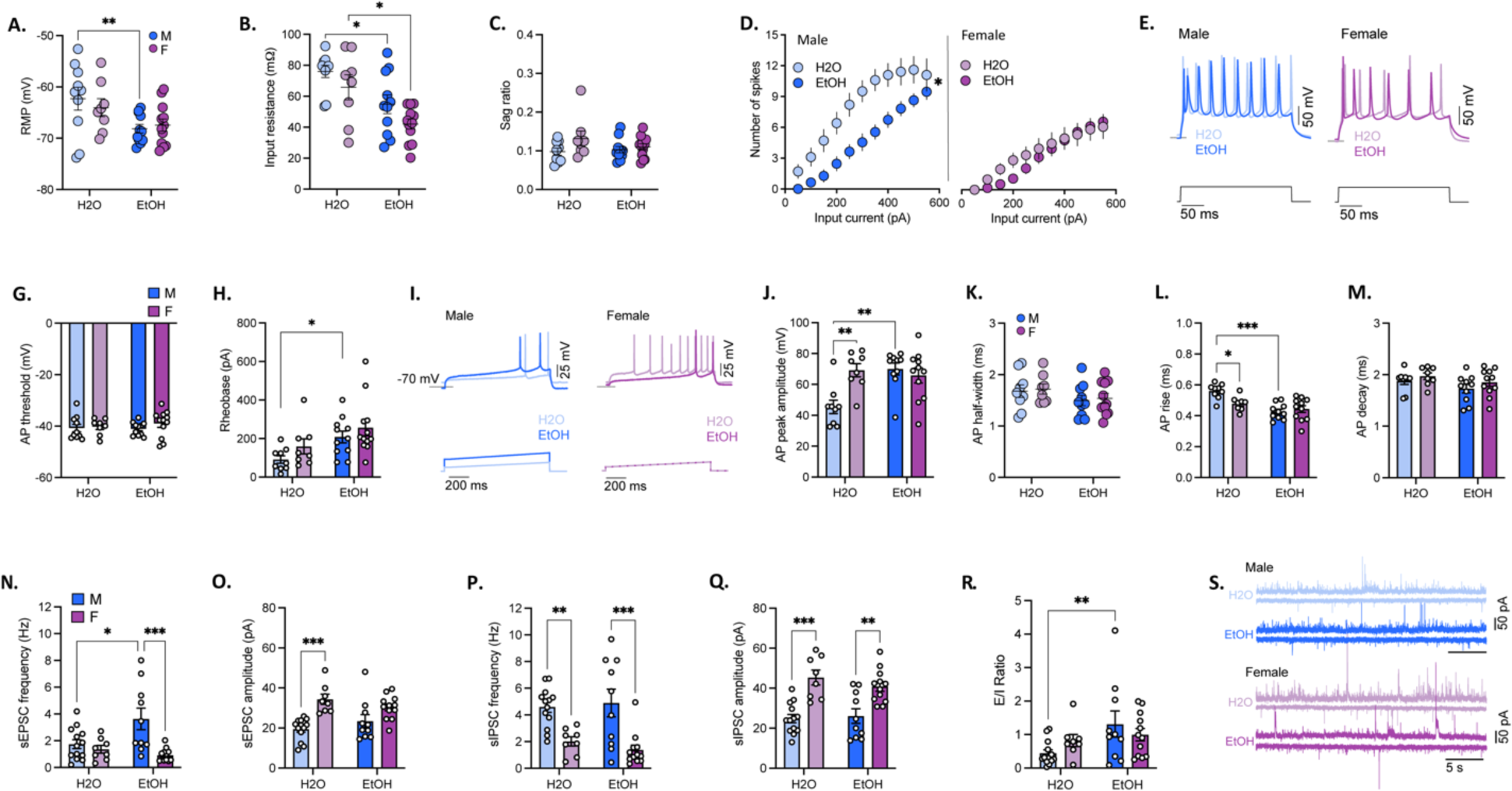
Insula Pdyn neurons are differentially altered by intermittent alcohol access in male and female mice. Whole-cell patch clamp recordings were obtained from Pdyn neurons in the insula of male and female Pdyn^GFP^ mice following 8 wks of intermittent alcohol (EtOH) access or water control (H2O) to assess alcohol’s effects of alcohol the (A-C) intrinsic properties, (D-M) excitability (at RMP), and (N-S) synaptic transmission of insula Pdyn neurons. (A) alcohol increased the resting membrane potential (RMP) of Pdyn neurons in female mice but not male mice, and had no effect on (B) input resistance or (C) sag ratio in either sex. (D-I) In male but not female mice, alcohol decreased the excitability of Pdyn neurons as indicated by (D) decreased firing to increasing current steps and (H) increased rheobase (minimum current needed to elicit firing). (G) There was no significant effect of alcohol on Pdyn action potential (AP) threshold. (E, I) Representative traces from male (left, blue) and female (purple, right) mice show (E) total firing elicited by a 600 pA current step, and (I) the amount of current required to fire an action potential (rheobase) determined using a 120 pA/1 s ramp protocol. There was no effect of alcohol on the action potential (AP) threshold or any measures of AP kinetics assessed, including (J) amplitude, (K) half-life, (L) rise time, or (M) decay time. There was a main effect of sex on (M) AP decay time, as Pdyn neurons of female mice exhibited shorter decays times. (N-S) Alcohol did not affect synaptic transmission in Pdyn neurons in either sex aside from producing a (Q) decrease in sIPSC amplitude in female but not male mice. There was no difference between EtOH and H2O groups in (N) sEPSC frequency, (O) sEPSC amplitude, or (P) sIPSC frequency in male and female mice. Sex differences in spontaneous synaptic transmission in Pdyn neurons were observed, as female mice exhibited (O) higher sEPSC amplitude, (P) lower sIPSC frequency and amplitude, and higher E/I ratio as compared to male mice and (H) EtOH-exposure increased the ratio of excitatory to inhibitory input onto Pdyn neurons. (S) Representative traces of sIPSC (top trace) and sEPSC (bottom trace) obtained from individual Pdyn neurons in male and female water (light blue and purple, respectively) and alcohol (dark blue and purple, respectively) exposed mice. * p < 0.05, ** p < 0.01, # p < 0.05 main effect of sex.

Next, we evaluated the impact of IA on synaptic transmission in Pdyn neurons (Fig. 5F-K). We found no effect of IA or sex on sEPSC frequency (males: n = 9 cells/4 mice EtOH and H2O; females: n = 11 cells/5 mice EtOH, n = 9 cells/4 mice H2O; [sex x treatment interaction: *p* = 0.401; sex main effect: *p* = 0.280; treatment main effect: *p* = 0.260]), suggesting that the frequency of glutamatergic synaptic events is similar in male and female mice and not altered by long-term IA. When we assessed GABAergic transmission, we found that females received less inhibitory input onto insula Pdyn neurons than male mice, as supported by a main effect of sex on sIPSC frequency [*F*_(1,34)_ = 4.75, *p* = 0.036]. As with sEPSCs, we found no effect of IA on sIPSC frequency [sex x treatment interaction: *p* = 0.819; treatment main effect: *F*_(1,36)_ = 4.71, *p* = 0.159]. Interestingly, when we assessed the ratio of excitatory to inhibitory (E/I) input onto insula Pdyn neurons, we found significant main effects of sex [*F*_(1,34)_ = 5.08, *p* = 0.031] and treatment [*F*_(1,34)_ = 6.10, *p* = 0.018] but no group x sex interaction [*p* = 0.754]. This finding indicates that there is greater excitatory synaptic drive in Pdyn neurons of male mice, and that IA increases E/I in this neuronal population in both males and females. When event amplitude was assessed, we found that sE/IPSC amplitudes were lower in male Pdyn neurons as compared to females, as indicated by a significant main effects of sex [sEPSC: *F*_(1,34)_ = 18.5, *p* < 0.001; sIPSC: *F*_(1,34)_ = 12.1, p < 0.001]. Whereas there was no effect of IA on sEPSC amplitude in male and female mice [sex x treatment: *p* = 0.802; treatment main effect: *p* = 0.222], sIPSC amplitude was decreased in IA exposed animals, as revealed by a significant main effect of treatment [*F*_(1,34)_ = 4.23, *p* = 0.047] but no interaction of sex x treatment [*p* = 0.330].

We subsequently examined the effect of long-term IA on intrinsic properties, excitability, and synaptic transmission in insula DLPNs of male mice (n = 11 cells/4 mice EtOH, n = 10 cells/4 mice H2O) and female mice (n = 12 cells/5 mice EtOH, n = 8 cells/4 mice), as show in in Fig. 6. In both male and female mice, we found that some intrinsic properties of insula DLPN were altered by IA (Fig. 6A-C). Specifically, 8 weeks of IA decreased RMP (Fig. 6A) in male but not female mice [IA vs H2O: males, *t*_37_ = 2.75, *p* = 0.009; females, *p* = 0.135; treatment x sex interaction: *p* = 0.431; treatment main effect: F_(1,37)_ = 9.03, p = 0.005; sex main effect: *p* = 0.737] and input resistance (Fig. 6B) in male and female mice [IA vs H2O: males, *t*_37_ = 2.88, *p* = 0.013; females, *t*_37_ = 3.09, *p* = 0.008; treatment x sex interaction: *p* = 0.812; treatment main effect: *F*_(1,37)_ = 17.8, *p* < 0.001; sex main effect: *F*_(1,37)_ = 4.67, *p* = 0.037]. There was no effect of IA on sag ratio (Fig. 6C) in either sex [treatment x sex interaction: *p* = 0.228; treatment main effect, *p* = 0.460; sex main effect, *p* = 0.069]. Long-term IA also reduced the excitability of insula DLPN (Fig. 6D-I) in male mice as we found a significant decrease in the number of action potentials fired at increasing current steps (Fig. 6D) in males [treatment x current interaction: *F*_(10,190)_ = 2.92, *p* = 0.002; treatment main effect: *F*_(1,19)_ = 10.4, *p* = 0.005; current main effect: *F*_(10,190)_ = 92.0, *p* < 0.001], but not in females [treatment x current interaction: *p* = 0.052; treatment main effect: *p* = 0.557; current main effect: F_(10,170)_ = 62.3, *p* < 0.001]. This was further supported by an increase in rheobase (Fig. 6H) in IA exposed male but not female mice [IA vs H2O: males, *t*_19_ = 2.43, *p* = 0.020; females, *p* = 0.059; sex x treatment interaction: *p* = 0.788; treatment main effect: *F*_(1,37)_ = 9.53, *p* = 0.004; sex main effect: *p* = 0.103]. There was no effect of IA on AP threshold (Fig. 6G) [Fs < 1 for treatment x sex interaction, and main effects of treatment and sex main]. Representative traces of insula DLPN firing to a 600 pA current step (Fig. 6E) and a 120 pA/s ramp protocol (Fig. 6I) in male (left, blue) and female (right, purple) H2O and IA mice. Additionally, we found several effects of IA on action potential kinetics in male mice, including an effect of IA on AP peak amplitude (Fig. 6J) [treatment x sex interaction: F_(1,34)_ = 9.81, p = 0.004; main effects of treatment: F_(1,34)_ = 5.65, *p* = 0.023, and sex: F_(1,34)_ = 4.83, *p* = 0.035], and on AP rise time (Fig. 6L) [treatment x sex interaction: F_(1,34)_ = 6.98, p = 0.012; treatment main effect: F_(1,34)_ = 20.7, *p* < 0.001; and sex main effect: *p* = 0.094]. Follow-up analyses showed that in male mice only IA produced a significant increase in AP peak amplitude [males: t_34_ = 3.92, *p* < 0.001; females: *p* > 0.999 ] and a significant decrease in AP rise time [males: t_34_ = 5.11, *p* < 0.001; females: *p* = 0.378] as compared to H2O controls. Analyses also revealed that on both measures there were significant sex difference in H2O control animals, as AP peak amplitude was lower [t_34_ = 3.58, p = 0.002] and rise time higher [t_34_ = 2.94, p = 0.012] in H2O males compared to females. There were no effects of IA or sex on AP half-width (Fig 6K) [Fs < 1 for treatment x sex interaction and sex main effect; treatment main effect: p = 0.095], or decay (Fig 6M) [treatment x sex interaction: F < 1; treatment main effect: p = 0.094; sex main effect: p = 0.175].

**Figure 6.**
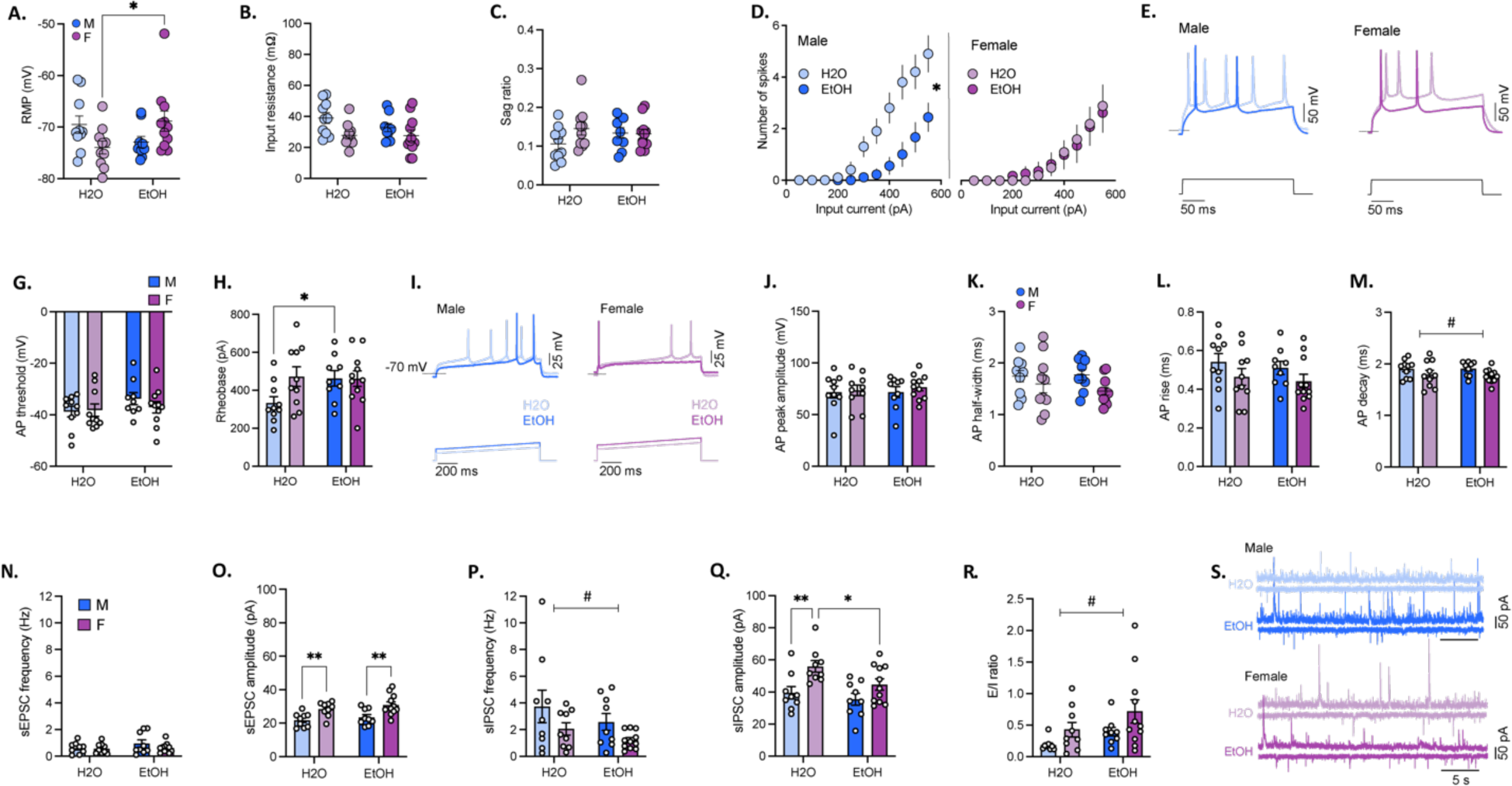
Insula deep layer pyramidal neurons (DLPNs) are differentially altered by intermittent alcohol access in male and female mice. Whole-cell patch clamp recordings were obtained from insula deep layer pyramidal neurons (DLPN) in male and female mice following 8 wks of intermittent alcohol (EtOH) access or water control (H2O). Alcohol’s effects of on the (A-C) intrinsic properties, (D-M) excitability (at RMP), and (N-S) synaptic transmission of insula DLPN were assessed. (A-B) Intrinsic properties of insula DLPN were altered by alcohol, as it produced a (A) decrease in DLPN resting membrane potential (RMP) in male mice, and (B) decrease in input resistance in males and females. (C) There was no effect of alcohol on DLPN sag ratio. (D-I) Alcohol decreased DLPN excitability in male but not female mice, as indicated by (D) decreased firing to increasing current steps and (H) increased rheobase (minimum current needed to elicit firing). G) There was no significant effect of alcohol on DLPN action potential (AP) threshold. (E, I) Representative traces from males (left, blue) and females (purple, right) show (E) total firing elicited by a 600 pA current step, and (I) the amount of current required to fire an action potential (rheobase) determined using a 120 pA/1 s ramp protocol. In male mice, alcohol altered AP kinetics of insula DLPN, as it (J) increased AP peak amplitude and (L) decreased AP rise time. There was no effect of alcohol in female mice on these measures. In H2O controls, APs elicited by insula DLPN in female mice had a (J) higher peak amplitude and (L) shorter rise time as compared to male mice. There was no effect of alcohol and no sex differences found for (K) AP half-width or (M) decay time. (N-S) Alcohol produced sex-specific effects on synaptic transmission in insula DLPN. (N) Alcohol increased sEPSC frequency but not amplitude in DLPN of male mice and had no effect in female mice. Excitatory events were lower in frequency in EtOH females and higher in amplitude in H2O females as compared to males within each group. Alcohol had no effect on sIPSC (P) frequency or (Q) amplitude in either sex. In female mice, inhibitory events were lower in frequency and higher in amplitude compared to males. (R) Alcohol increased the ratio of excitatory to inhibitory (E/I) events in male mice, and there was no difference in E/I ratio by sex. (S) Representative traces of sIPSC (top trace) and sEPSC (bottom trace) obtained from individual DLPN in the insula of male and female water (light blue and purple, respectively) and alcohol (dark blue and purple, respectively) exposed mice. * p < 0.05, ** p < 0.01, *** p < 0.001.

Synaptic transmission was next assessed in DLPNs of IA-exposed and water control male and female mice (Fig. 6F-K). In these experiments we found that long-term IA produced a sex-dependent increase in excitatory synaptic transmission. This was revealed by a significant sex x treatment interaction on sEPSC frequency (males: n = 10 cells/5 mice EtOH, n = 13 cells/4 mice H2O; females: n = 12 cells/5 mice EtOH, n = 8 cells/4 mice H2O; [*F*_(1,39)_ = 6.60, *p* = 0.014]) as well as a significant main effect of sex [*F*_(1,39)_ = 11.7, *p* = 0.001] but not group [*p* = 0.117]. Post-hoc analyses revealed that IA increased DLPN sEPSC frequency in male mice [*p* = 0.023] but not female mice [*p* > 0.999], which subsequently resulted in a significant increase in sEPSC frequency in IA-exposed males compared to females [*p* < 0.001]. Conversely, there was no effect of long-term alcohol intake on inhibitory synaptic transmission in insula DLPN neurons, as sIPSC frequency did not differ between IA mice and water controls. This finding was demonstrated by the absence of both a sex x treatment interaction [*p* = 0.442] and main effect of treatment [*p* = 0.776]. Analyses yielded a main effect of sex on sIPSC frequency [*F*_(1,39)_ = 25.3, *p* < 0.001] and post-hoc comparisons revealed that H2O and IA exposed males exhibited higher sIPSC frequency as compared to females of the same treatment group [H2O: *p* = 0.006; IA: *p* < 0.001]. We next assessed E/I ratio in insula DLPNs and found that IA significantly increased synaptic drive in this neuronal population. This was demonstrated by a significant main effect of treatment [*F*_(1,39)_ = 5.13, *p* = 0.029] but not sex [*p* = 0.881] and no sex x treatment interaction [*p* = 0.141]. Follow-up analyses showed that there was a significant difference in E/I ratio between IA and water exposed males [*p* = 0.008] but not females [*p* = 0.608], indicating that IA increased excitatory synaptic drive in male mice only. Moreover, we found sex-related differences in event amplitude, as sEPSC and sIPSs were higher in female mice. This was illustrated by a significant main effect of sex on sE/IPSC amplitude [sEPSC: *F*_(1,39)_ = 23.6, *p* < 0.001; sIPSC: *F*_(1,39)_ = 36.5, *p* < 0.001] and post-hoc analyses showing significant differences in sEPSC between male and female H2O mice [*p* < 0.001] and sIPSC between male and female H2O [*p* < 0.001] and IA mice [*p* = 0.004].

## Discussion

In this study we used converging pharmacological and genetic approaches to gain insight into how DYN/KOR signaling in the insula contributes to excessive alcohol consumption in male and female mice. In addition, we used slice electrophysiology to probe physiological changes in insula neuronal populations, focusing on DYN expressing neurons and Layer 5 pyramidal neurons. Across these studies we identified numerous sex differences at the behavioral and physiological level. This adds to a growing literature demonstrating sex differences in how KOR systems can modulate behavior and neuronal function.

### Divergence between Pharmacology and Genetic Approaches

In this study, there were several key differences between the site-directed pharmacological (blocking KOR) and genetic approaches (deleting KOR/Pdyn) we used to probe the impact on drinking. In male mice, we saw an early impact on alcohol consumption that diminished over time. We hypothesized that could be due to the waning effects of nor-BNI or a shift to a different state that is insensitive to local KOR antagonism. To test this, we then infused nor-BNI in another set of male mice that had already escalated their alcohol intake, and found no effect on consumption. This suggested that in males only, local KOR antagonism in the insula could reduce early alcohol consumption, but following escalation, it was no longer sensitive. This result was surprising, as in many other studies, KOR antagonist effects emerge as animals become dependent or escalate their consumption (Walker et al., 2011; Erikson et al., 2018). However, it is notable that KOR antagonists can impact drinking in more acute models of escalated consumption, such as Drinking in the Dark (DID) (Anderson et al., 2019; Haun et al., 2020). The antagonist approach does have limitations, including difficulty to accurately quantify spread and potential off-target actions of the compound.

To address these limitations, we used a genetic approach, similar to our previous work in the central nucleus of the amygdala (CeA) (Bloodgood et al., 2021). In contrast to local inhibition, we found that deletion of KOR had no effect on any measures of alcohol intake. This divergence is likely due to the dissociation between the receptors that the local antagonist can block compared to the target population of KOR ablated by a conditional deletion approach. It is likely that the insula receives inputs that presynaptically express KOR on their terminals, and this population of KOR can be blocked by local pharmacology but not by deletion of KOR from insula neurons. Another possibility is that deletion of KOR from insula neurons also leads to removal of presynaptically-expressed KOR in regions downstream from the insula. Thus, in these output regions, KOR signaling may play a distinct role in alcohol consumption that differs from that of the insula. Finally, we cannot exclude the possibility that some of the differences between approaches may be due to issues related to knockout efficiency and viral expression/spread.

We also examined how deletion of insula Pdyn could impact alcohol drinking and found that it led to a robust reduction of alcohol consumption in both males and females, with some sex-related differences. Specifically, in female mice there was a reduction in alcohol intake in both early and late phases of the IA procedure. In males, the reduction in alcohol intake was only apparent in the later phases of IA, where mice typically escalate their consumption. Deletion of Pdyn in females also reduced sucrose intake, suggesting that insula Pdyn may more broadly regulate consumption of rewarding solutions in female mice. This is especially interesting as recent studies have implicated the insula in feeding related behaviors (Stern et al., 2021). However, the experiments in Stern et al. (2021) were performed in male mice and implicated a distinct population of central amygdala-projecting insula neurons in feeding and satiety signaling. Thus, we are cautious to speculate whether our observations may extend to other signaling systems or neuronal populations within the insula. Interestingly, we found that Pdyn and KOR manipulations differentially impacted alcohol consumption, specifically in terms of the phase of intake that was impacted by our manipulations. One possible reason for this may be that when we delete Pdyn, we are altering both local and distal neuropeptide release. There are a number of insula targets that could contribute to this effect, such as the CeA, which has been implicated in KOR regulation of alcohol consumption. Further, deletion of pDyn may have additionally impacted other peptides, such as leu-enkephalin, which is derived from precursor molecules pDyn and pro-enkephalin (Akil et al., 1984; Evans et al., 1988). Thus, in using this genetic approach, we cannot exclude the possibility that other peptides may have contributed to our findings. Another potential caveat to this work is that the mouse lines we have used to perform these genetic studies are not fully backcrossed on to a C57BL/6J background. We observed similar differences in baseline consumption in an earlier study, investigating the role of Dyn/KOR in the CeA (Bloodgood et al., 2021). In the future, CRISPR based tools may provide a means to circumvent this concern.

Notably, none of the manipulations we performed that altered alcohol consumption altered avoidance behavior, assessed via the elevated plus maze. This suggests that the role of KOR/DYN system may be distinct from regulation of avoidance behavior. Previous work has shown that the insula and its output to the extended amygdala are engaged by and contribute to negative affective states in mice (Centanni et al., 2019; Luchsinger et al., 2021). However, these experiments examined the role of insula during the distinct state of forced abstinence from alcohol or during an explicit behavior, specifically struggling during restraint stress. Although we did not explore insula DYN/KOR during forced abstinence or restraint stress, this would be an interesting direction for future work, and may further illuminate the impact of alcohol-induced DYN/KOR signaling shifts in affective outcomes. Additional work from the Hopf and Lasek groups has shown that the insula is associated with aversion-resistant and compulsive-like alcohol intake (Seif et al., 2013; Chen et al., 2015; Chen and Lasek, 2020; Darevsky and Hopf, 2020; De Oliveira Sergio et al., 2021). We did not explore that phenotype in our model, and it represents another path for exploration.

### Alcohol consumption has complex effects in distinct insula cell types

Given that Pdyn deletion decreased alcohol intake, we next wanted to evaluate the impact of long-term alcohol consumption on the properties of insula Pdyn neurons. We found that with the excitability of these neurons, there were divergent effects depending on sex. In males, we found a reduction in current-evoked firing and an increase in rheobase, consistent with reduced excitability. In female mice, we found that IA increased the resting membrane potential in Pdyn neurons only. There was a main effect of alcohol drinking on the E/I ratio across both sexes, however there was reduced basal GABAergic tone in insula Pdyn neurons from females, suggesting possible differences in the function of insula interneurons or external GABA inputs on insula Pdyn neurons. Taken together, the results suggest that IA induces long-term synaptic plasticity in insula Pdyn neurons in males and are possibly indicative of a synaptic scaling process. This is especially interesting considering recent results suggesting synaptic scaling plays a role in learning in the closely related gustatory cortex (Wu et al., 2021). The increase in RMP and synaptic balance in insula Pdyn neurons from female mice could be linked to the greater effect of DYN deletion in females, however our data do not explicitly address this possibility. It is noteworthy that these findings, in particular the sex differences, are similar to our recent study exploring the impact of binge-like alcohol drinking on DYN neurons in the CeA (Bloodgood et al., 2021), and align with recent data from the Sparta lab demonstrating sex-specific effects of alcohol on insula neuronal plasticity (Marino et al., 2020). This raises the important issue that sex is a critical factor to consider when evaluating the insula and KOR ligands in preclinical models, as has been noted by Chartoff and Mavrikaki (Chartoff and Mavrikaki, 2015).

We also evaluated the impact of long-term alcohol consumption on Layer 5 pyramidal neurons in the IC, as outputs from the insula have been implicated in a range of complex cognitive and physiological processes (Gogolla, 2017), and found to be altered following alcohol exposure (McGinnis et al., 2020). A similar pattern emerged with these layer 5 neurons showing reductions in excitability, with an increase in E/I ratio from males, which is suggestive of a scaling-like phenomena, but with no differences in the properties of layer 5 insula from female mice. This further demonstrates that alcohol drinking induces a sex-dependent plasticity and highlights the need for additional work to resolve how this may impact the behaviors we observed. Previous studies from the Besheer lab (Jaramillo et al., 2018b, 2018a, 2018c) and the Hopf lab (Seif et al., 2013) have shown that an insula to nucleus accumbens pathway is important for alcohol self-administration and aversion-resistant alcohol drinking. This may indicate one possible mechanism for how the cellular changes we observed may promote increased alcohol consumption. The Hopf lab also found a role for insula outputs to brainstem in punished alcohol drinking but not alcohol only drinking (De Oliveira Sergio et al., 2021), suggesting that while this path is important, it may not be related to the drinking phenotypes seen. However, there is also intriguing work that insula outputs to the BLA can play a critical role in learning (Yiannakas et al., 2021). This is especially exciting when taken with recent data supporting the role of the BLA in alcohol reinforcement (Faccidomo et al., 2021). Future studies should focus on investigating plasticity in these distinct outputs, as it can provide important insight into how alcohol can impact insular cortex circuits.

As summarized in our working model in Fig. 7, our combined data suggest that KOR/DYN signaling in the insula can regulate alcohol consumption and there are key sex differences in the mechanism. Pharmacology suggests that in males, there is an early KOR signal in the insula that plays a role in induction but not maintenance of escalated alcohol drinking. In contrast, Pdyn deletion can reduce alcohol consumption during the later phases of drinking, suggesting a shift to KOR sensitivity in downstream structures, potentially the CeA. In females, Pdyn deletion appears to robustly impact drinking in both the early and late of IA. Given that sucrose intake was concomitantly decreased, it is possible that Pdyn deletion may more generally affect consummatory or appetitive responses in female mice. There are similar sex-dependent changes on neuronal function, however our data does not specifically identify how they are related to escalated consumption. Notably, our findings underscore the importance of accounting for sex as a biological variable in alcohol-related behavior, and specifically the mechanisms by which KOR regulates alcohol consumption. As a potential target for treatment of AUD, it is important to identify these mechanisms and determine how they relate to sex differences in clinical populations.

**Figure 7.**
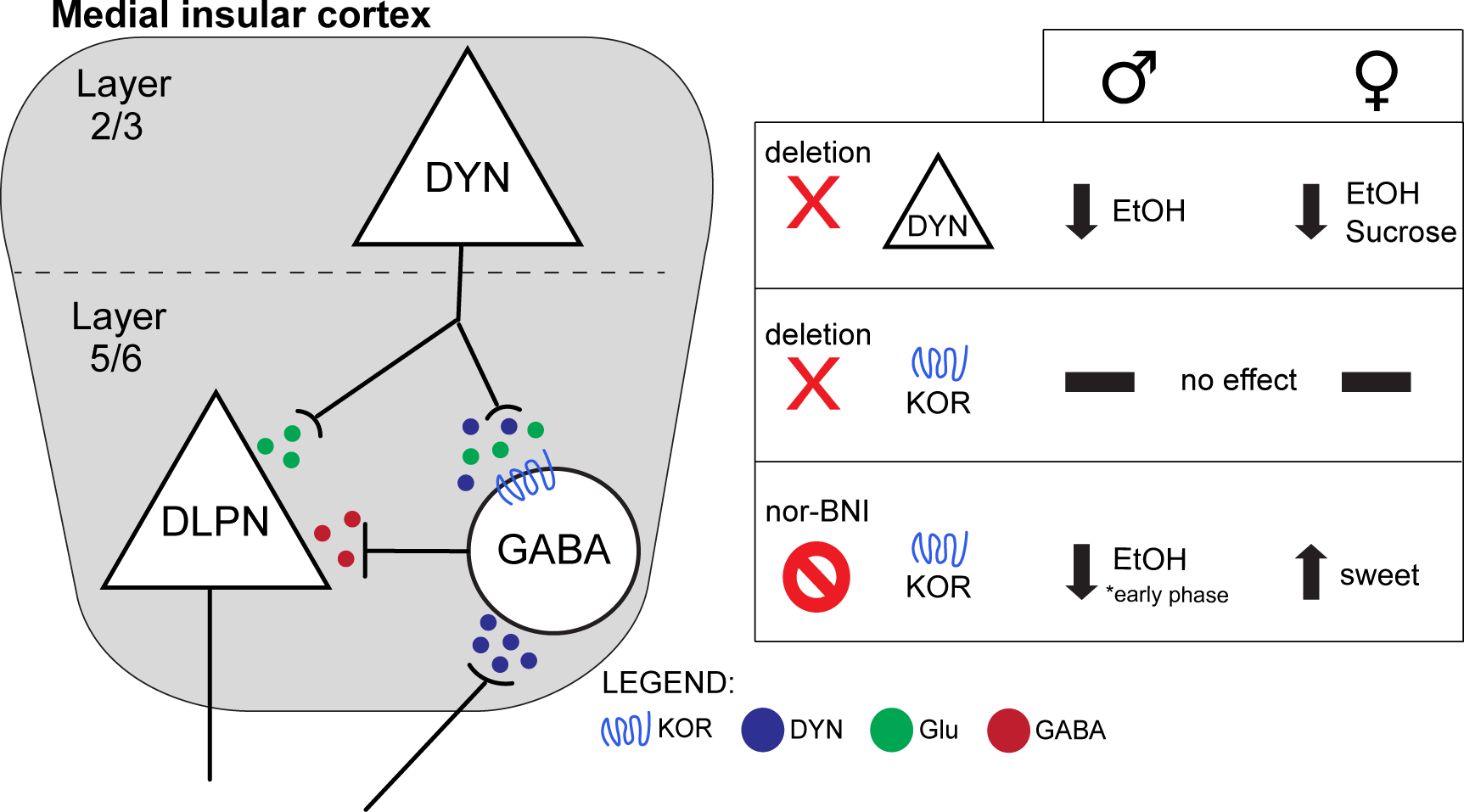
Working model of insular cortex DYN/KOR involvement in drinking behavior in male and female mice. Summary of experimental outcomes highlighting how different manipulations of KOR signaling can impact behavior.

## Acknowledgements

This work was supported by National Institutes of Health grants P60 AA011605 (TLK/MN), F32 AA026485 (MMP), K99/R00 AA028298 (MMP), U01 AA020911 (TLK), R01 AA025582 (TLK), R01 AA025582-S1 (TLK/MMP)

---

**Figure S2.**
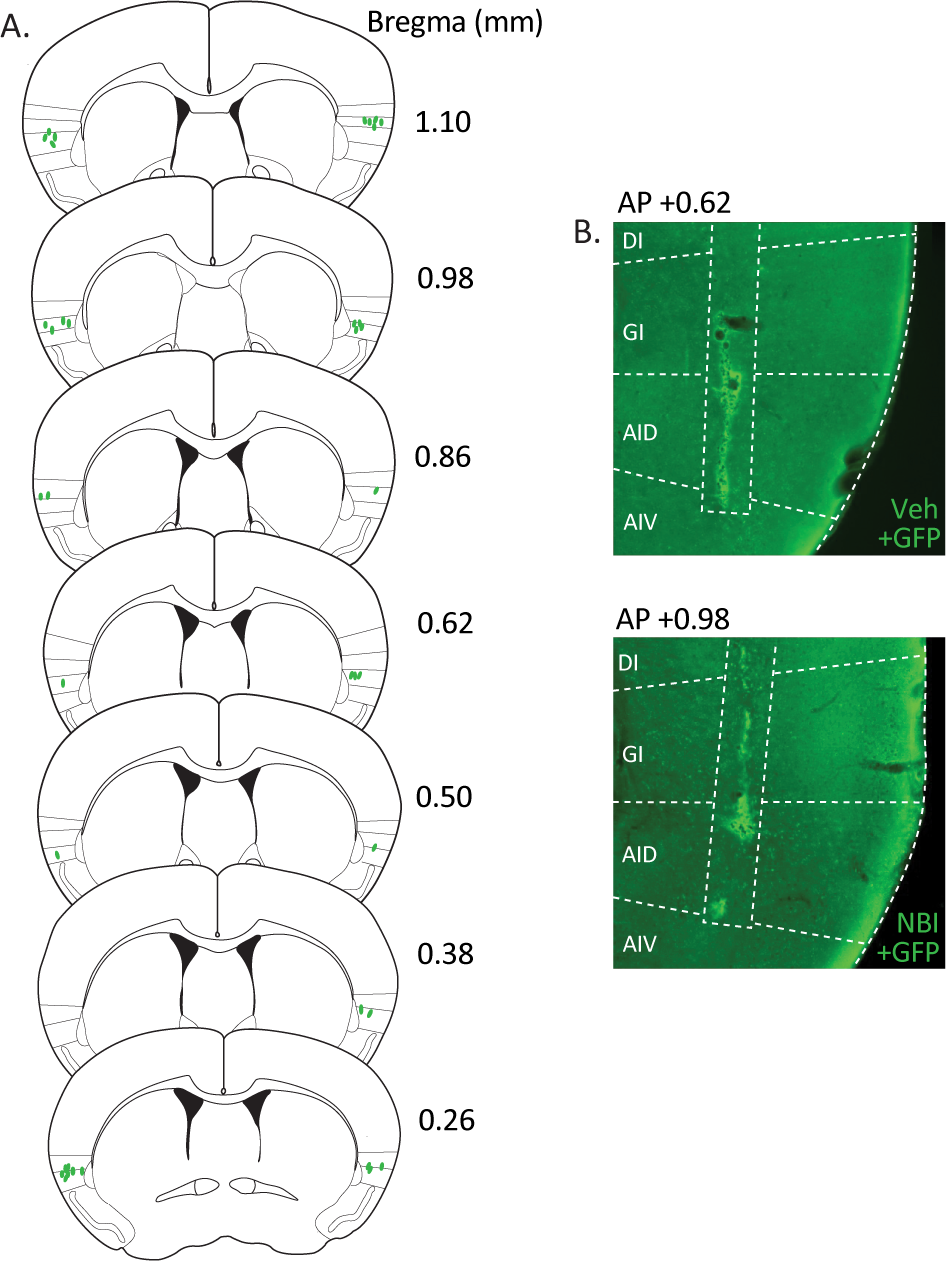
Location of intracranial drug infusions. (A) Reconstruction of injector tip placements for nor-BNI and Veh treated female mice shown in Fig. 2 that were co-injected with 50 nL of an AAV encoding for GFP. Numbers indicate distance from bregma (in mm) for each coronal section. (B) Representative micrograph showing the location of the injector tract (rectangular dashed outline) and its most ventral point of termination, which was used to map the injector tip placement illustrated in (A) for Veh (top) and NBI (bottom) treated mice. Numbers indicate the location of each slice in the anterior-poster (AP) plane (in mm from bregma). Dashed lines segment each division of the insula shown. Insular cortex divisions: DI, dysgranular; GI, granular; AID, dorsal agranular; AIV, ventral agranular.

**Figure S3.**
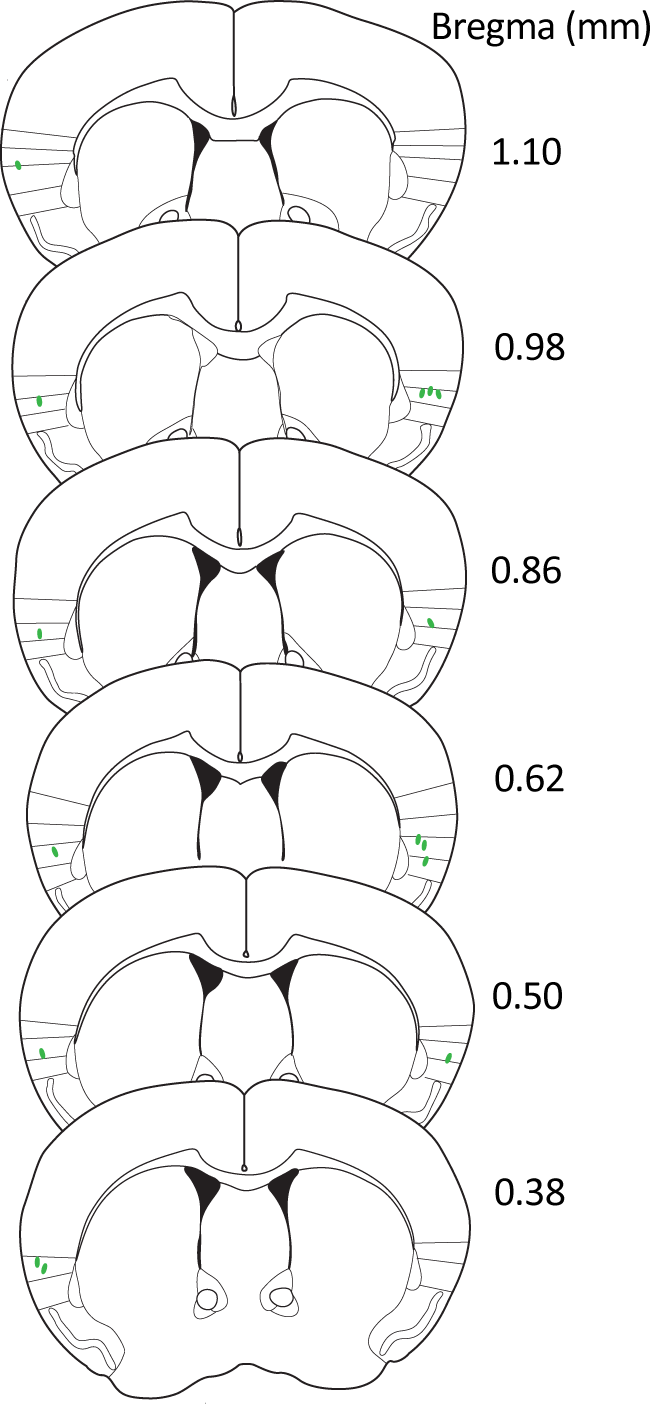
Location of intracranial drug infusions. Reconstruction of injector tip placements for nor-BNI and Veh treated female mice shown in Fig. 3 that were co-injected with 50 nL of an AAV encoding for GFP. Numbers indicate distance from bregma (in mm) for each coronal section. Figure S3. Location of intracranial drug infusions. Reconstruction of injector tip placements for nor-BNI and Veh treated female mice shown in Fig. 3 that were co-injected with 50 nL of an AAV encoding for GFP. Numbers indicate distance from bregma (in mm) for each coronal section.

